# Non-sedating benzodiazepines cause contractile paralysis and tissue damage in the parasitic blood fluke *Schistosoma mansoni*

**DOI:** 10.1101/694588

**Authors:** Paul McCusker, Yeunus Mian, Guanguan Li, Michael D. Olp, V. V. N. Phani Babu Tiruveedhula, Farjana Rashid, Lalit Kumar Golani, Brian C. Smith, James M. Cook, John D. Chan

**Affiliations:** Department of Cell Biology, Neurobiology & Anatomy, Medical College of Wisconsin, Milwaukee, WI 533226, USA; Department of Chemistry and Biochemistry, Milwaukee Institute of Drug Discovery, University of Wisconsin-Milwaukee, Milwaukee, WI 53211, USA; Department of Biochemistry, Medical College of Wisconsin, Milwaukee, WI 533226, USA

## Abstract

Parasitic flatworm infections (e.g. tapeworms and fluke worms) are treated by a limited number of drugs. In most cases, control is reliant upon praziquantel (PZQ) monotherapy. However, PZQ is ineffective against sexually immature parasites, and there have also been several concerning reports of cestode and trematode infections with poor PZQ cure-rates, emphasizing the need for alternative therapies to treat these infections. We have revisited a series of benzodiazepines, given the known anti-schistosomal activity of meclonazepam (MCLZ). MCLZ was discovered in the 1970’s but was not brought to market due to dose-limiting sedative side effects. However, in the decades since there have been advances in our understanding of the benzodiazepine GABA_A_ receptor sub-types that drive sedation and the development of sub-type selective, non-sedating ligands. Additionally, the sequencing of flatworm genomes reveals that parasitic trematodes and cestodes have lost GABA_A_R-like ligand gated anion channels, indicating that MCLZ’s anti-parasitic target is likely distinct from the human receptors that drive sedation. Therefore, we screened a library of classical and non-sedating 1,4-benzodiazepines against *Schistosoma mansoni* and identified a series of imidazobenzodiazepines that immobilize worms *in vitro*. One of these hits, Xhe-II-048 also disrupted the parasite tegument, causing extensive vacuole formation beneath the apical membrane. The imidazobenzodiazepine compound series identified has a dramatically lower (∼1 log) affinity for human central benzodiazepine binding site and is a promising starting point for the development of novel anti-schistosomal benzodiazepines with minimal host side-effects.

**Author Summary:** Over 200 million people are infected with schistosomiasis, yet there are limited therapeutic options available to treat this disease. The benzodiazepine meclonazepam is known to cure both intestinal and urinary schistosomiasis in animal and human studies, but dose-limiting sedation has been a barrier to its development. Little is known about the structure-activity relationship of meclonazepam and other benzodiazepines on schistosomes, or the identity of the parasite receptor for these compounds. However, schistosomes lack obvious homologs to the human GABA_A_Rs that cause sedation. This indicates that the parasite target of this drug is distinct from the host receptors that underpin dose-limiting side effects of meclonazepam, and raises the possibility that benzodiazepines with poor GABA_A_R affinity may still retain anti-parasitic effects. Here, we report an *in vitro* screen of various benzodiazepines against schistosomes, and the identification of hit compounds that are active against worms yet possess reduced affinity for the human GABA_A_Rs that cause sedation.

## 1. Introduction

Over 200 million people are infected with the parasitic blood flukes that cause the neglected tropical disease schistosomiasis (1). Over 90% of infections occur in sub-Saharan Africa, where the disease kills 300,000 persons / year (2). The pathology associated with chronic infection adds to the disease burden, putting the socioeconomic cost of schistosomiasis (70 million disability adjusted life years) near HIV/AIDs, malaria or tuberculosis (1, 3). However, despite these enormous costs treatment relies on just one broad-spectrum drug, praziquantel (PZQ) (4). PZQ treatment has high cure rates of 70-90% (5, 6), but it is concerning that a subset of infections in human and animal populations appear to be refractory to treatment (7-9), either due to PZQ’s lack of efficacy against recently acquired, immature parasites worms (10, 11) or standing genetic variation in parasite populations. The latter possibility is especially concerning in regard to the potential emergence of PZQ-resistant parasites, and consideration needs given to whether PZQ-monotherapy will be sufficient to achieve schistosomiasis elimination (12).

One lead compound with proven anti-schistosomal activity is the benzodiazepine meclonazepam ((S)-3-methylclonazepam, MCLZ). MCLZ was discovered in the 1970’s and found to cure both the mature and immature parasites that cause urinary and intestinal forms of schistosomiasis (13). Development of this lead stalled in the 1980’s due to dose-limiting sedation in human trials (14-17). However, we have revisited benzodiazepines as potential anti-parasitic leads given advances in two areas. First, it is now understood that the sedative effects of benzodiazepines are driven by α1-subunit containing GABA_A_Rs (18). This has enabled the design of various benzodiazepines with reduced affinity towards α1-containing GABA_A_Rs to treat various conditions involving GABA_A_Rs that contain other α subunits such as asthma and schizophrenia (19). Second, with recent advances in the sequencing of parasitic helminth genomes there is an abundance of data available to establish whether flatworm parasites possess GABA_A_Rs (20). If GABA_A_Rs are not present in parasitic worms, then it is possible that the structure-activity requirements of anti-parasitic compounds may differ significantly from those mediating benzodiazepine binding to host GABA_A_Rs, offering the opportunity to develop ligands with increased parasite selectivity. Here, we have profiled the repertoire of *S. mansoni* ligand gated ion channels and, having found no obvious parasite GABA_A_Rs, screened a library of benzodiazepines to identify compounds that display anti-parasitic activity and exhibit reduced mammalian GABA_A_R affinity.

## 2. Materials and Methods

### Bioinformatic prediction of flatworm Ligand-Gated Ion Channels

Putative ligand-gated ion channels were curated from a diversity of organisms sampling vertebrates (human), arthropods (*Drosophila melanogaster*), nematodes (*Caenorhabditis elegans*), and mollusks (*Aplysia californica*). The predicted proteomes of these organisms were searched for gene products containing a ligand-gated ion channel (LGIC) Pfam domain (PF02932). These were used to generate hidden Markov probability models (HMMER v3.2.1), which were used to search the predicted proteomes of various free-living *(Schmidtea mediterranea, Macrostomum lignano*) and parasitic (*Schistosoma mansoni, Echinococcus multilocularis*) flatworms for putative LGICs. Resulting candidates were filtered based on number of predicted transmembrane domains (TOPCONS) (21) and manually inspected for the presence of a characteristic Cys loop F/YPxD motif. The retained sequences were then aligned (MUSCLE) and degapped (GapStreeze v2.1.0, 50% tolerance) to enable construction of Maximum Likelihood phylogenetic trees (LG model with 500 bootstrap replicates).

### Chemicals

A complete list of chemical structures is provided in Supplemental Table 2. Compounds were either sourced commercially (Toronto Research Chemicals (Meclonazepam) and Sigma Aldrich (clonazepam, nitrazepam, diazepam, bromazepam, flurazepam, lorazepam, flunitrazepam)) or synthesized by the Cook Lab as reported in references (22-25). Structures were generated in ChemDraw (v17.1) and clustered by physiochemical properties using ChemMine (26).

### Adult schistosome mobility assays

Female Swiss Webster mice infected with *S. mansoni* cercariae (NMRI strain) were sacrificed 49 days post infection by CO_2_ euthanasia. Adult schistosomes were recovered by dissection of the mesenteric vasculature. Harvested schistosomes were washed in DMEM (ThermoFisher cat. # 11995123) supplemented with HEPES (25mM), 5% heat inactivated FCS (Sigma Aldrich cat. # 12133C) and Penicillin-Streptomycin (100 units/mL). Worms were cultured in 6 well dishes (4-5 male worms per well) in the presence of various test compounds or DMSO vehicle control overnight (37°C / 5% CO_2_). Worms were imaged the next day to record movement phenotypes using a Zeiss Discovery v20 stereomicroscope and a QiCAM 12-bit cooled color CCD camera controlled by Metamorph imaging software (version 7.8.1.0). 1 minute recordings were acquired at 4 frames per second and saved as a .TIFF stack, which was imported into ImageJ for analysis. An outline of the workflow used to quantify movement from these video recordings is shown in Supplemental Figure 1. Maximum intensity projections were generated for the entire stack of images (241 frames for a 1-minute recording) and integrated pixel values were measured for the resulting composite image, allowing normalization of movement relative to DMSO control treated worms. Inhibition of movement IC_50_ values were calculated using GraphPad Prism v8.1.1 and are expressed ± 95% confidence intervals. Data represents mean ± standard error for ≥3 independent experiments. Significance (*) was determined by unpaired t-test at a threshold of 0.05. Animal work was carried out with the oversight and approval of the Laboratory Animal Resources facility at the Medical College of Wisconsin. All animal experiments followed ethical regulations approved by the Medical College of Wisconsin IACUC committee.

**Figure 1.**
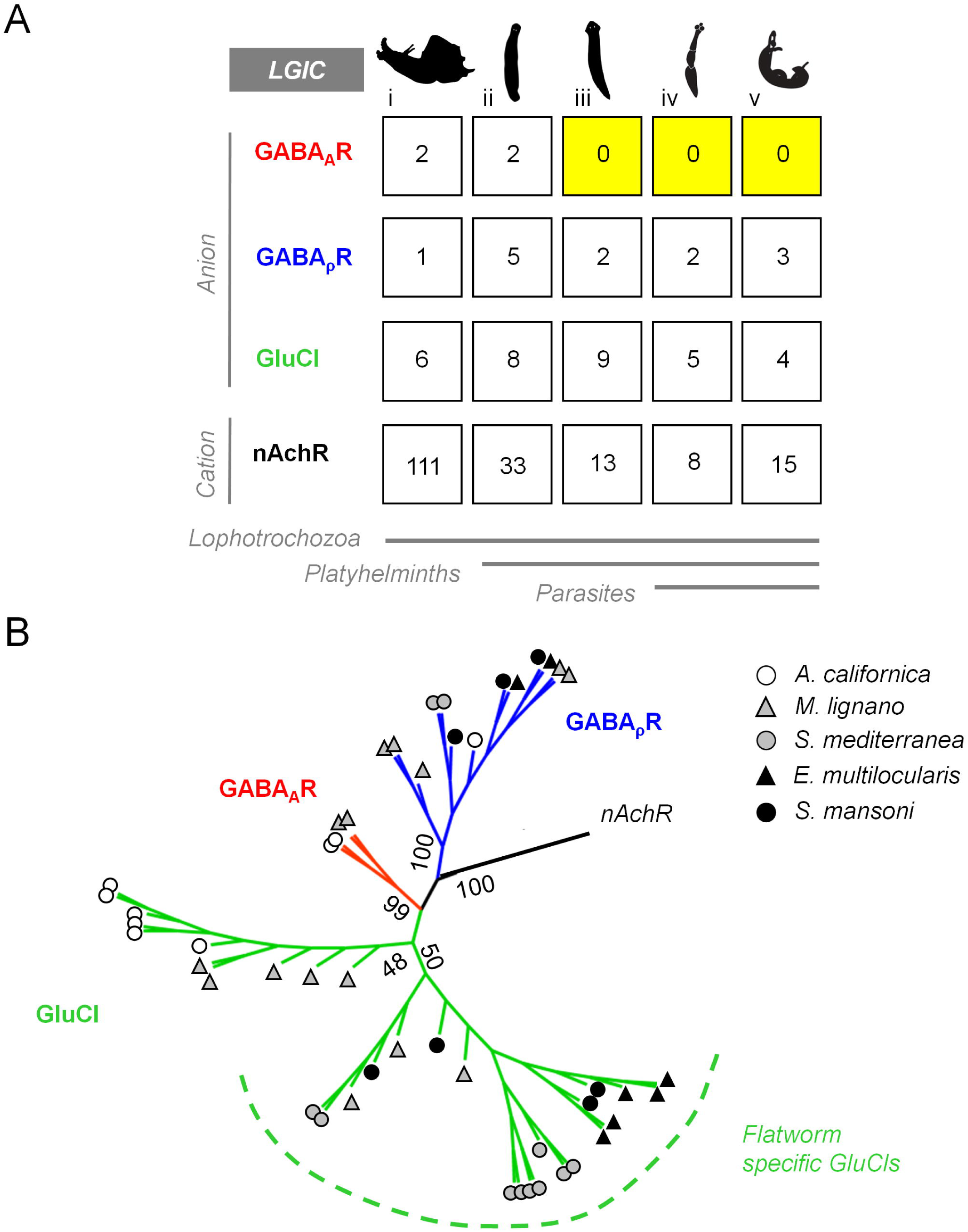
Comparison of free-living and parasitic flatworm Cys-Loop Ligand Gated Ion Channels (LGICs). **(A)** Cys-loop LGICs were curated from the genomes of a non-flatworm lophotrochozoan (i. *A. californica*) as well as free-living (ii. *M. lignano*, iii. *S. mediterranea*) and parasitic flatworms (iv. *E. multilocularis*, v. *S mansoni*) and clustered into various families of GABA_A_R-like, GABA_ρ_R-like, GluCl-like and nAchR-like sequences. **(B)** Cladogram of flatworm and lophotrochozoan LGICs, with a clade of GABA_A_R subunits (red), GABA_ρ_R subunits (blue) and two distinct clades of GluCl subunits (green) that include a previously reported flatworm-specific group of receptors (31). Bootstrap percentage is shown at nodes (500 replicates).

### Transmission Electron Microscopy

Adult worms were harvested and recovered as above. Fixation was carried out overnight at 4°C in 2.5% glutaraldehyde/2% paraformaldehyde in 0.1 M sodium cacodylate (pH 7.3). Worms were washed 3 × 10 minutes in 0.1 M sodium cacodylate and post-fixed for 2 hours on ice in reduced 1% osmium tetraoxide. Worms were then 2 × 10 minutes in distilled water and stained overnight at 4°C in alcoholic Uranyl Acetate. Worms were rinsed in distilled water, dehydrated in 50%, 75% and 95% MeOH, followed by successive 10 minute rinses in 100% MeOH and acetonitrile. Worms were incubated in a 1:1 mix of acetonitrile and epoxy resin for 1 hour prior to 2 × 1 hour incubations in epoxy resin. Worms were then cut transversely and embedded overnight in epoxy (60°C). Ultra-thin sections (70 nm) were cut onto bare 200-mesh copper grids and stained in aqueous lead citrate for 1 minute and sections were imaged on a Hitachi H-600 electron microscope fitted with a Hamamatsu C4742-95 digital camera) operating at an accelerating voltage of 75 kV.

### Binding Assays

Binding assays for benzodiazepines against mammalian GABA_A_Rs were performed, measuring displacement of [3H] flunitrazepam (0.4 nM) from crude brain membrane preparations. Rat cerebral cortex membrane homogenate (80 µg protein) was incubated with 0.4 nM [3H]-flunitrazepam in 50 mM Tris-HCl (pH 7.7), plus test compounds (screened at concentrations ranging from 0.1 nM to 10 µM), and non-specific binding was assessed by incubation with diazepam (3 µM). Following incubation for 60 min at 4°C, samples were vacuum filtered through glass fiber filters (GF/B, Packard) presoaked with 0.3% PEI and rinsed with ice-cold 50 mM Tris-HCl (Unifilter, Packard). Filters were dried and counted for radioactivity in a scintillation counter (Topcount, Packard). *K*_*i*_ values of test compounds were calculated by the Cheng-Prusoff equation.

### GABA_A_R Modeling

Benzodiazepines were docked to the human GABA_A_R cryo-EM structure (PDB ID 6HUO)(27) using Schrödinger Maestro suite v2019-1. 3-D ligand structures were generated using the LigPrep module. Water molecules further than 5 Å from the protein surface were removed, hydrogen bonds were optimized at pH 7.0 and the structure was minimized in the OPLS3e forcefield. Ligands were docked into a 10× 10×10 Å grid centered on the bound alprazolam molecule in the GABA_A_R structure using Glide (28) in ExtraPrecision (XP) mode, and output poses were ranked by XP GlideScore. Figures were generated using PyMOL 2.3.0.

## 3. Results

### 3.1 GABA_A_Rs are not the schistosome targets of meclonazepam

The benzodiazepine meclonazepam (MCLZ) is an effective anti-schistosomal drug, but the sedative side effects of MCLZ coincide with the anti-parasitic dose (14). The sedative side effects of MCLZ are likely driven by human GABA_A_Rs – specifically those heteromeric receptors that contain the α1 subunit (18). These receptors account for approximately 60% – 80% of brain GABA_A_Rs (18, 29). However, the parasite target of MCLZ remains unknown. Since the earliest reports of MCLZ’s anti-schistosomal activity it has been noted that these effects are not replicated by other benzodiazepines that also bind GABA_A_Rs with high affinity (13, 30). Therefore, we considered that the parasite target of MCLZ may be distinct from GABA_A_Rs, since it is not even clear whether flatworms possess this class of ligand-gated ion channel (LGIC) (31, 32).

In order to comprehensively search the repertoire of flatworm LGICs, we generated hidden Markov probability models (HMMER v3.2.1) using LGICs curated from a diversity of organisms (humans, *D. melanogaster, C. elegans, A. californica*). These were used to search the predicted proteomes of various free-living *(S. mediterranea, M. lignano*) and parasitic (*S. mansoni, E. multilocularis*) flatworms to retrieve putative LGICs. Candidates were manually inspected for accurate number of predicted transmembrane domains and the presence of a characteristic Cys-loop F/YPxD motif. The resulting sequences clustered into four groups corresponding to three groups of ligand gated anion channels (Glutamate-gated Chloride Channels (GluCl), GABA_A_Rs and GABA_ρ_Rs) and one group of ligand gated cation channel-like gene products (nicotinic acetylcholine receptor (nAchR)-like) (Supplemental Table 1). Previously reported schistosome GluCls (31) and cholinergic receptors (33) clustered alongside sequences consistent with their functional characterization. GABA_A_Rs are clearly present in the mollusk *A. californica* and free-living flatworm *M. lignano*, but they appear to have been lost in free-living planarians and the parasitic flatworms *E. multilocularis* and *S. mansoni* (Figure 1).

### 3.2 Non-sedating imidazobenzodiazepines cause parasite contractile paralysis

If schistosomes lack GABA_A_Rs, then the parasite target of MCLZ may have different structural requirements for ligand binding than the MCLZ – GABA_A_R interaction that drives sedation. If so, then it should be possible to identify parasite-selective benzodiazepines that retain anti-schistosomal activity but lack affinity at mammalian GABA_A_Rs that contain α1-subunits. Therefore, we screened a library of compounds that included various α1GABA_A_R-sparing compounds against adult male *S. mansoni* and assessed action on schistosomes *in vitro*. Worms were harvested from mice 7-weeks post-infection and cultured in test compound (30 µM) overnight, after which video recordings were acquired to measure drug effects on worm movement relative to vehicle negative control (DMSO 0.1%) and MCLZ (5 µM) positive control. This primary screen of 180 compounds identified 19 ligands that phenocopied MCLZ, exhibiting a coiled, contractile phenotype and paralysis (Figure 2A). Active compounds were then re-screened at 10 µM to refine the hits to the most active compounds. This resulted in the prioritization of two chemical series. The first were MCLZ derivatives, including clonazepam (CLZ, lacking the C3 methyl group of MCLZ), and the derivative MYM-I-91A with the phenyl C2’ halogen substituted from a chlorine to a fluorine (Figure 2A&B). The second series was a group of imidazobenzodiazepines (SH-I-055, XliHeII-048 and XHE-II-048) that also varied in the C3 and phenyl C2’ positions (Figure 2A&C).

**Figure 2.**
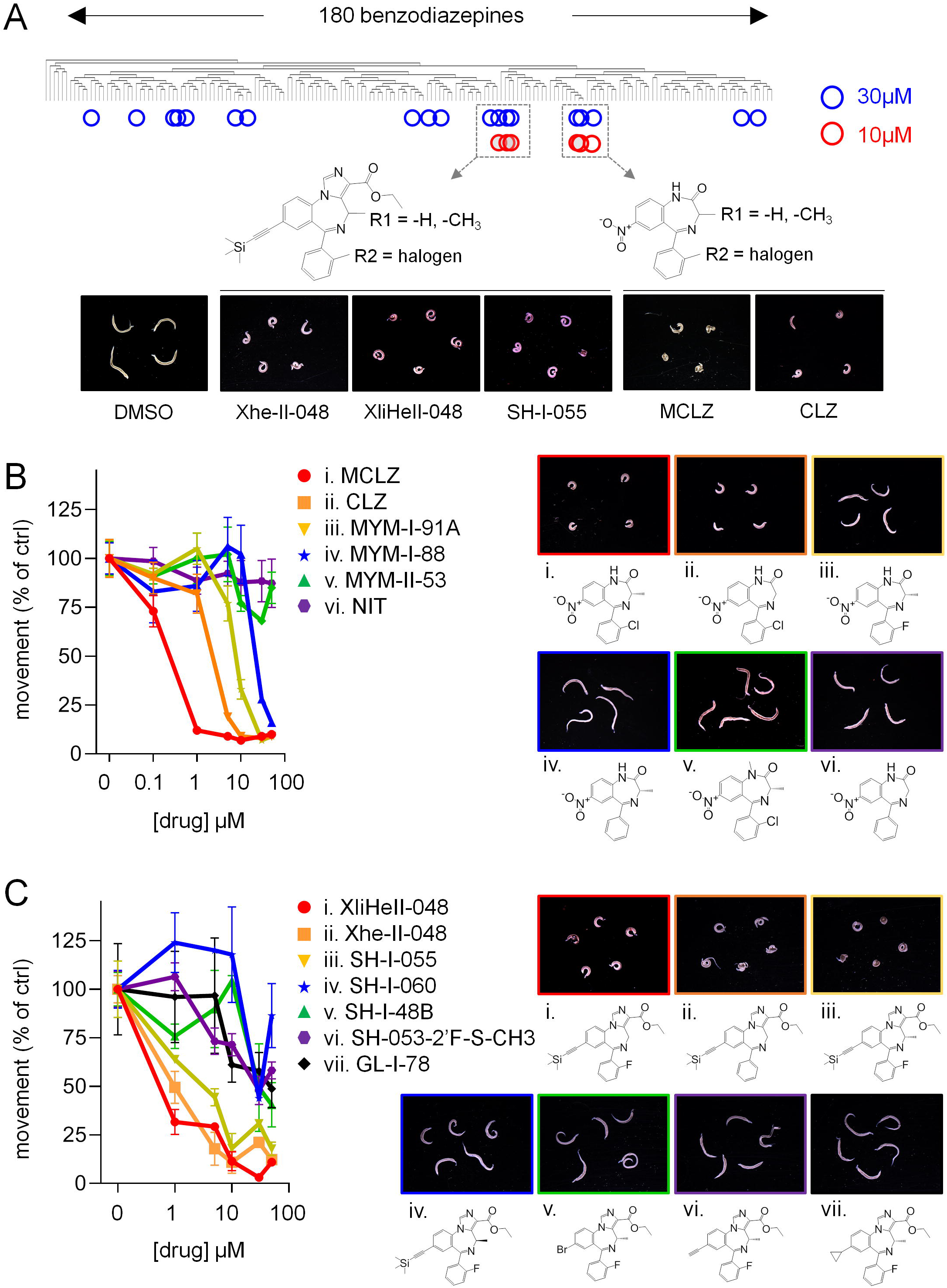
Identification of benzodiazepines with *in vitro* activity against *S. mansoni*. **(A)** 180 benzodiazepines were screened for ability to contract and paralyze schistosomes *in vitro*. Compounds were initially screened at 30 µM (hits = blue circles), and active compounds were re-screened at 10 µM (hits = red circles). This prioritized imidazobenzodiazepines with a TMS-acetylene moiety (left) and meclonazepam-like compounds (right). Structure-activity relationship of **(B)** a series of MCLZ derivatives and **(C)** a series of imidazobenzodiazepines. MCLZ = meclonazepam, CLZ = clonazepam, NIT = nitrazepam. Left = movement dose response curves for parasites exposed to each compound. Right = Images of drug treated worms (10 µM, overnight).

The structure activity relationship of these two series was explored, assessing the comparative potency of various derivatives at impairing schistosome movement. Qualitative observations on the activity of various MCLZ-like benzodiazepines have been previously reported (13, 30, 34), and the activity of the MCLZ-like series was largely consistent with these studies. The rank order of potency was MCLZ (IC_50_ 160 nM), CLZ (IC_50_ 1.9 µM), MYM-I-91A (IC_50_ 7.2 µM), MYM-I-88 (IC_50_ 26.0 µM), followed by MYM-II-53 and nitrazepam which were inactive at concentrations as high as 50 µM (Figure 2B). The structure-activity relationship of the imidazobenzodiazepines was explored, assessing the ability of the three hit compounds and structurally related, less active compounds to evoke coiled, contractile paralysis (Figure 2C). The most active compounds, XliHeII-048 (IC_50_ 540 nM) and Xhe-II-048 (IC_50_ 850 nM), contain an imidazole-ester group at the benzodiazepine N1-C2 position and a trimethylsilyl (TMS) acetylene group at the C7 position. The two compounds differ only by the addition of a fluorine at the phenyl C2’ position on XliHeII-048. A third compound, SH-I-055 (IC_50_ 1.4 µM) was identical to XliHeII-048 except for the addition of a chiral (S)-methyl group at the C3 position. When this chiral methyl group was in the (R) orientation there was a marked decrease in potency (compound SH-I-060). Finally, compounds retaining SH-I-055’s (S)-methyl group but with varying C7 modifications in place of the TMS acetylene group all showed dramatically decreased affinity (GL-I-78, SH-I-48B and SH-053-2’F-S-CH3 with a cyclopropyl, bromine and alkyne group, respectively).

The potency of XliHeII-048 and Xhe-II-048 at inhibiting worm movement was comparable to the active ligands in the MCLZ derivative series with IC_50_ values in the high nanomolar range (Figure 2B&C). Therefore, binding assays were performed to compare the relative affinities of these two chemical series for mammalian GABA_A_Rs (Figure 3). As expected, MCLZ potently displaced [3H]-flunitrazepam from rat brain membrane preparations (*K*_*i*_ 2.4 nM). The related compound CLZ bound GABA_A_Rs with an even higher affinity (*K*_*i*_ 0.82 nM). Analogs MYM-I-91A and MYM-II-53 also displayed high affinity (*K*_*i*_ 12.1 nM and 4.1 nM, respectively). MYM-I-88, which lacks a halogen on the phenyl ring, displayed markedly reduced binding (*K*_*i*_ 76.4 nM). The imidazobenzodiazepine series displayed a GABA_A_R affinity roughly one log less potent than MCLZ (Xhe-II-048 *K*_*i*_ = 2.5 µM, SH-I-055 *K*_*i*_ = 1.7 µM, XliHeII-048 *K*_*i*_ = 1.6 µM).

**Figure 3.**
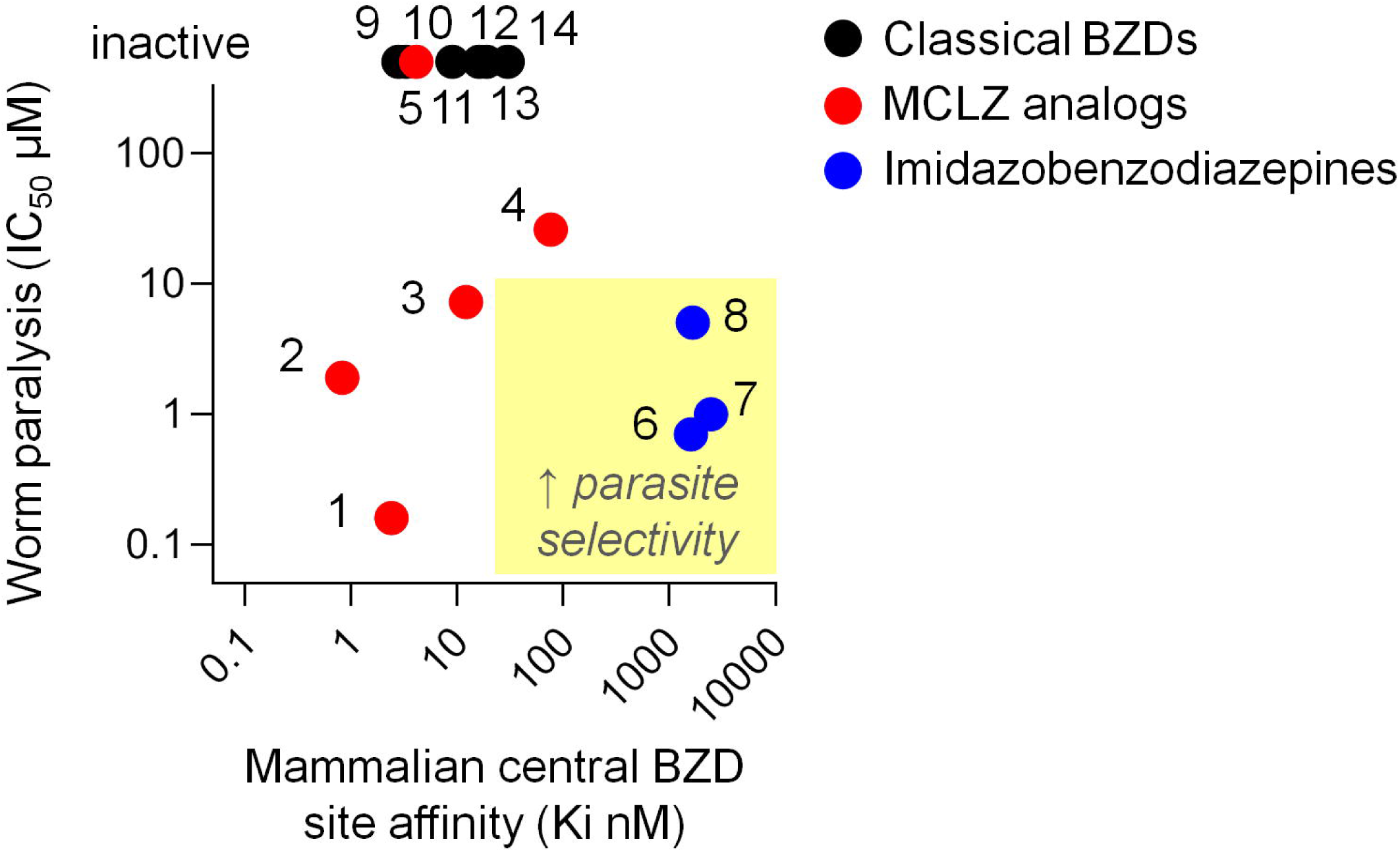
Relative mammalian GABA_A_R affinity and schistosome potency of various benzodiazepines. Scatter plot of mammalian central benzodiazepine binding site affinity (*K*_*i*_) verses schistosome activity (movement IC_50_). Sedating compounds active against worms (i.e. MCLZ) fall within the lower left quadrant. Desired compounds that lack sedation but retain anti-parasitic effects fall within the lower right quadrant. 1 = MCLZ, 2 = CLZ, 3 = MYM-I-91A, 4 = MYM-I-88, 5 = MYM-II-53, 6 = Xhe-II-048, 7 = XliHeII-048, 8 = SH-I-055, 9 = flunitrazepam, 10 = lorazepam, 11 = diazepam, 12 = flurazepam, 13 = nitrazepam, 14 = bromazepam. Mammalian central benzodiazepine binding site *K*_*i*_ values for compounds 9-14 are from reference (51).

### 3.3 Imidazobenzodiazepine Xhe-II-048 causes structural damage to parasite tissue

Given the parasite-selectivity of imidazobenzodiazepines (Figure 3, blue) relative to MCLZ-like compounds (Figure 3, red), we investigated the effects of XliHeII-048, Xhe-II-048 and SH-I-055 on schistosome tissues in more detail. Specifically, we were interested in drug-evoked damage to the parasite tegument, which is a feature of many anti-schistosomal compounds (35). Worms treated with DMSO vehicle control, MCLZ (5 µM) or various imidazobenzodiazepines (10µM) overnight, fixed and processed for imaging by transmission electron microcopy (TEM). Imaging transverse schistosome cross sections revealed a typical body wall structure in DMSO treated worms, with alternating layers of schistosome muscle, followed by the tegument basal membrane, tegument syncytium, and tegument apical membrane. In MCLZ treated worms, tissue layers exhibit are disrupted, with pervasive vacuolization of the tegument (Figure 4A). The tegument of Xhe-II-048 treated worms displayed a similar pattern of extensive vacuole distribution beneath the apical membrane, while worms treated with XliHeII-048 and the less potent imidazobenzodiazepine SH-I-055 displayed normal tissue ultrastructure (Figure 4A).

**Figure 4.**
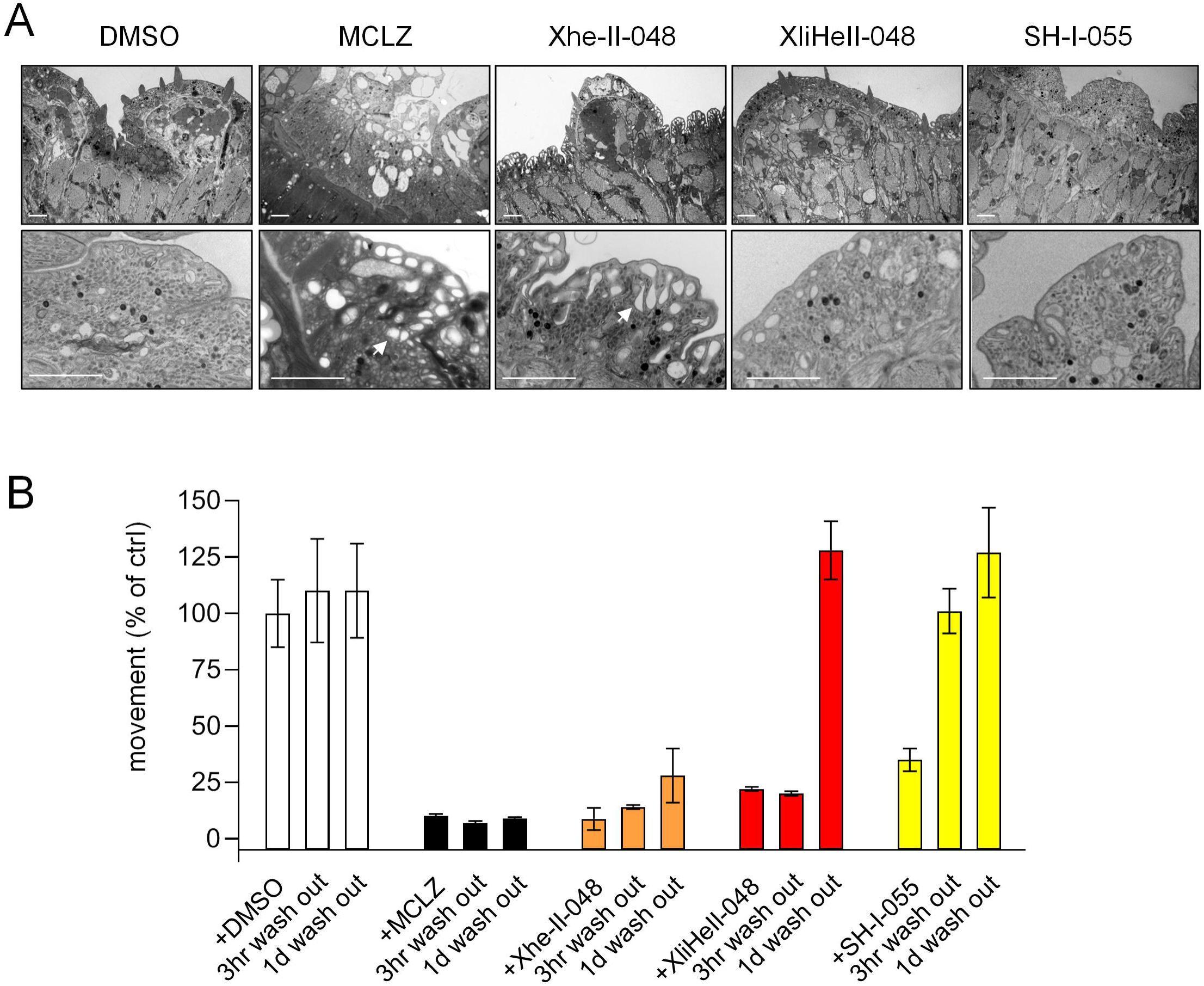
Xhe-II-048 damages the schistosome tegument. **(A)** Transmission electron microscopy images of transverse sections of *S. mansoni* exposed to either DMSO control, MCLZ (5 µM) or various imidazobenzodiazepines (10 µM, 14 hours). Dorsal tegument is oriented to the top. Scale = 2 µm. Arrowed = vacuoles beneath the tegument apical membrane. **(B)** Movement of worms treated with DMSO, MCLZ, or TMS-acetylene imidazobenzodiazepine compounds (10 µM, 14 hours), and recovery at various timepoints (3 hours, 1 day) following drug washout.

MCLZ is a schistocidal compound, and worms do not recover movement following drug washout (Figure 4B). Similarly, Xhe-II-048 treated worms did not recover movement up to one day following drug washout, consistent with the extensive ultrastructural damage caused by this compound (Figure 4A). Compounds XliHeII-048 and SH-I-055, which did not cause pervasive tegument damage, evoked only a transient paralysis, recovering by 1 day after drug washout.

## 4. Discussion

The anti-schistosomal activity of the benzodiazepine meclonazepam (MCLZ) was discovered in the 1970’s, but development of this lead as a human therapy for schistosomiasis stalled due to sedative side effects that coincide with the dose of drug needed to clear infections (13, 14). This is expected, given the structural similarity of MCLZ to centrally acting benzodiazepines such as CLZ and NIT that are clinically used as anxiolytics and display well known sedating effects. Some attempts were made in the 1980’s to antagonize the sedative effects of MCLZ using the GABA_A_R antagonist flumazenil. While flumazenil did not impair the anti-schistosomal effect of MCLZ (36), pharmacokinetic differences between flumazenil (administered by IV due to poor bioavailability, <1 hour elimination half-life (37)) and MCLZ (orally bioavailable, half-life up to 80 hours (38)) precluded the development of an admixture as a viable non-sedating therapeutic approach (15, 39). Research on MCLZ as an anti-schistosomal lead has slowed over the past several decades. However, we have revisited this compound based on recent helminth genomic data indicating that parasitic flatworms lack GABA_A_Rs (Figure 1), advances in our understanding of mammalian GABA_A_R subtypes that account for sedative side effects (18, 40), and advances in the development of non-sedating benzodiazepines with selectivity towards various GABA_A_R sub-types (22-25).

### 4.1 Schistosome genomes lack GABA_A_Rs

Sequenced genomes of parasitic trematode (*S. mansoni*) and cestode (*E. multilocularis*) flatworms appear to lack GABA_A_Rs, the benzodiazepine targets that cause sedation in humans (Figure 1) (32, 41, 42). Gene loss is common with the evolution of parasitism (43, 44), although GABA_A_Rs may also have been lost in the free living flatworm lineage since *M. lignano* contains GABA_A_R-like sequences but planarians do not. Within the flatworm phylum, Macrostomum appear to have diverged basal to the lineage containing Tricladida (planaria) and Neodermata (parasitic flatworms) (45), and flatworm GABA_A_R loss may have occurred after this split.

If parasitic flatworms lack GABA_A_Rs, might MCLZ act on related cys-loop ligand-gated ion channels (GluCls and GABAρRs)? Experimental and bioinformatic evidence indicates that this is unlikely. MCLZ has been shown to be inactive against SmGluCl, a representative of the class of flatworm chloride channels most similar to GABA_A_Rs (31). Recent structural data resolving the interactions of classical benzodiazepines with human GABA_A_Rs provide an explanation for this (27). Specifically, the human α1His102 residue that interacts with the benzodiazepine C7 position at the interface of the GABA_A_R α1 and γ subunits is replaced by an arginine in the flatworm-specific GluCls. This position is important, since human α4 and α6 GABA_A_R subunits also contain an arginine in this position, and the larger sidechain likely sterically clashes with classical benzodiazepines to render GABA_A_Rs comprised of α4 and α6 subunits benzodiazepine-insensitive. In the case of each of the three schistosome GABAρRs, the α1His102 position contains a negatively charged aspartic acid. This switch from a positively charged histidine sidechain may oppose interactions with the electron-dense benzodiazepine C7 position. The inactivity of benzodiazepines on flatworm cys-loop LGICs is consistent with the observation that, aside from MCLZ, and to a lesser degree the structurally similar compound CLZ, benzodiazepines lack anti-schistosomal activity ((13), Figure 3, Supplemental Table 2). While the flatworm target of MCLZ remains unidentified, these findings support the hypothesis that the parasite target is distinct from the human GABA_A_Rs that account for dose-limiting sedation.

### 4.2 S. mansoni are paralyzed by a class of α1GABA_A_R-sparing benzodiazepines

While benzodiazepines are typically considered GABA_A_R ligands with anxiolytic or sedative properties, numerous α1GABA_A_R-sparing members of this class have been developed with indications as diverse as anti-asthma to anti-viral medications (24, 46-48). Several imidazobenzodiazepines with a C7 trimethylsilyl (TMS) acetylene group caused schistosome contractile paralysis *in vitro* at high nanomolar – low micromolar concentrations (Figure 2A&C) and display low GABA_A_R binding affinity (Figure 3). These compounds were originally synthesized as part of a chemical series exploring α2 / α3-selective benzodiazepines as potential anxiolytic, anticonvulsant and antinociceptive leads (23, 25). The differing GABA_A_R affinities of MCLZ and the imidazobenzodiazepine hits may be due the interaction of the MCLZ N1 position with the α1 S205, which is disrupted by the addition of the imidazole ring. Modification of this N1 is also observed in the non-sedating antiviral benzodiazepine BDAA (47), although smaller alkyl groups are likely tolerated, such as the methyl on diazepam and MYM-II-53. Additionally, the large TMS-acetylene group at the C7 position of the imidazobenzodiazepine hit series may not be tolerated within the GABA_A_R binding pocket, where the MCLZ C7 nitro group is predicted to interact with the γ N60 sidechain (Figure 5).

**Figure 5.**
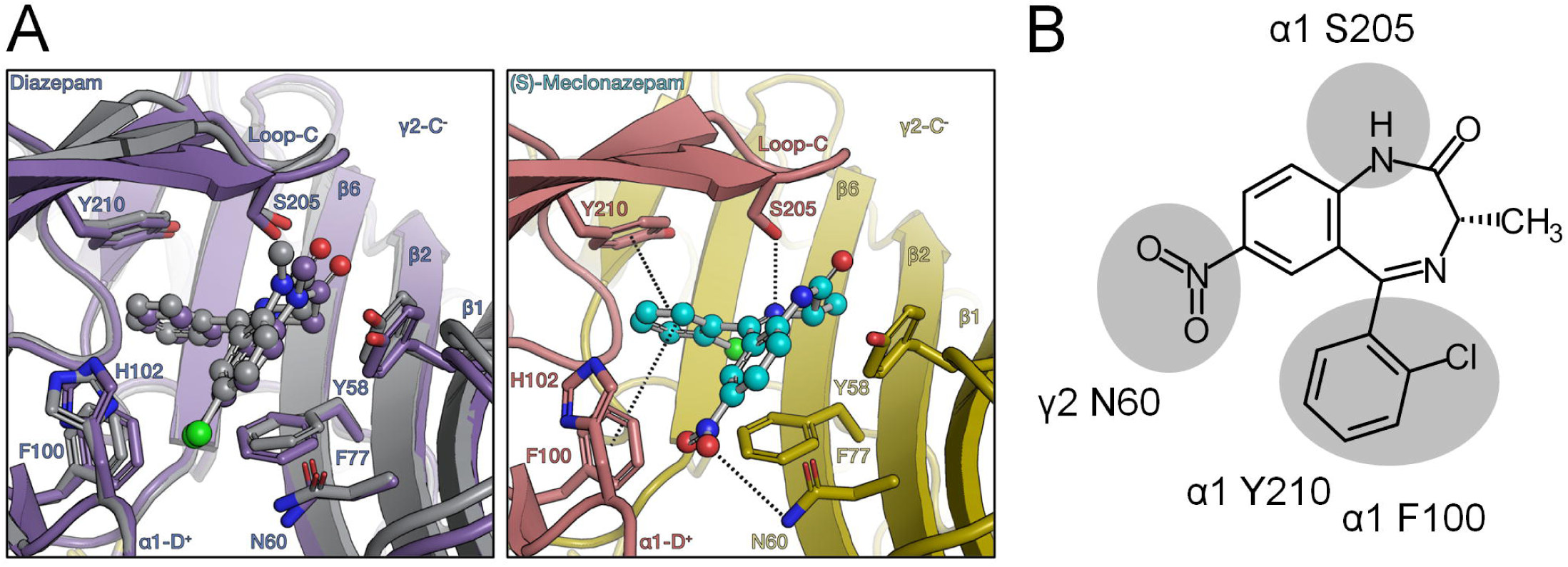
Modeling MCLZ interaction with host GABA_A_Rs. **(A)** *In silico* docking of meclonazepam to the benzodiazepine binding site of the solved human heteromeric α1β3γ2 GABA_A_R cryo-EM structure (27). Left = overlay of the diazepine bound cryo-EM structure (grey) and the docked diazepine ligand (purple), demonstrating that modeling procedures recapitulate experimentally observed ligand pose. Right = docked position of MCLZ, with predicted interactions with amino acid side chains on the α1 and γ2 subunits (dashed lines). **(B)** 2-D chemical structure of MCLZ, with potential GABA_A_R interactions highlighted.

While these ligands phenocopy MCLZ to a degree, it is unclear whether they act via the same schistosome receptor – or if they do bind the same receptor, whether they share a common binding pose. The target of MCLZ will need to be identified to generate hypothesis into ligand-receptor structure-activity relationships, as the structures of MCLZ and the TMS-acetylene imidazobenzodiazepines appear quite different. However, we can observe some similarities and differences in the SAR of the two series.

Two interesting positions are (i) the benzodiazepine C3 position, which is typically unmodified in classical benzodiazepines but contains a chiral methyl group in these two series, and (ii) the phenyl C2’ position, which is commonly halogenated in benzodiazepines with GABA_A_R affinity. The benzodiazepine C3 position is essential for MCLZ activity, the key difference between clonazepam and MCLZ is that clonazepam lacks a C3 methyl group. This results in a roughly 1 log decrease in potency (Figure 3). Chirality of this position is also important for 3-methylclonazepam, since the (R) enantiomer reportedly exhibits reduced anti-parasitic efficacy (34). On the other hand, C3 methylation decreased activity of XliHeII-048. The (S)-methylated compound SH-I-055 was ∼3 times less potent than XliHeII-048, and the (R)-methylated compound SH-I-060 was essentially inactive. This schistosome SAR is distinct from the binding of imidazobenzodiazepines to mammalian GABA_A_Rs, where (S) and (R) isomers have roughly equivalent binding affinities (49). The halogen on the phenyl C2’ position of MCLZ seems important for schistosome activity, given that nitrazepam (inactive against parasites) differs from clonazepam (movement IC_50_ 1.9 µM) in that the phenyl ring is unsubstituted. However, XliHeII-048 (which contains a fluorine at this position) and Xhe-II-048 (which possess an unsubstituted phenyl ring) appear equipotent (Figure 2C). In fact, the unsubstituted compound Xhe-II-048 was unique in evoking structural damage to the parasite tegument (Figure 4A).

Development of the Xhe-II-048 hit compound into a bona fide anti-schistosomal lead will likely require the modifications to improve metabolic stability *in vivo*. Specifically, the ester and TMS-acetylene groups will likely require substitution with bioisosteres. The imidazole ester group is likely rapidly hydrolyzed by carboxylesterases during first pass metabolism, and the TMS-acetylene group is also likely to be unstable *in vivo* (50). From the SAR shown in Figure 2C, is it apparent that loss of this group dramatically decreased potency. Nevertheless, this is the first report of non-sedating benzodiazepines screened against schistosomes. MCLZ has been shown to have anti-schistosomal activity in human clinical trials, but with an extremely narrow therapeutic index; the effective anti-parasitic dose (0.2 – 0.3 mg/kg) coincided with the dose at which sedation was reported (14). This work has identified benzodiazepine hits that exhibit potent anti-schistosomal effects *in vitro* and dramatically lower affinity for host α1GABA_A_Rs (Figure 3). Given the scarcity of new lead compounds to treat schistosomiasis, these data are valuable in advancing a pharmacophore that retains anti-schistosomal activity while displaying reduced sedation.

## Acknowledgements

This work was supported by funds from the Therapeutic Accelerator Program and a New Faculty Pilot Grant awarded to J.D.C by the Medical College of Wisconsin. We also gratefully acknowledge the Shimadazu Laboratory for Advanced and Applied Analytical Chemistry (UWM). Schistosome-infected mice were provided by the NIAID Schistosomiasis Resource Center at the Biomedical Research Institute (Rockville, MD) through NIH-NIAID Contract HHSN272201000005I for distribution via BEI Resources.

## Supporting Information

**S1 Fig. Quantification of schistosome movement.** Worm movement was quantified from video recordings (1 minute duration, 4 frames per second). **(A)** Video recordings in color (i) were converted to gray scale and inverted so that worms were transformed to dark silhouettes against a light background (ii). Video recordings (.tiff stacks of 241 images) were treated as a Z-stack, with a composite image of the maximum intensity from each frame integrated into one composite image (iii). **(B)** Movement was quantified by calculating the pixel intensity values of the drug treated composite and expressed relative to the DMSO vehicle control treated composite (left), producing a numerical quantification of movement across each dose (right).

**S1 Table. List of putative Aplysia and flatworm LGICs.** Sequence IDs reflect putative LGICs curated from *A. californica* (all NCBI deposited proteins for taxonomy ID 6500) *S. mansoni* (assembly v7) *E. multilocularis* (assembly EMULTI002), *S. mediterranea* (assembly SmedGD_c1.3) and *M. lignano* (assembly Mlig_3_7) and clustered into either nAch, GluCl, GABA_A_ or GABA_ρ_ like receptors. Other than *A. californica*, all assemblies were accessed via WormBase ParaSite.

**S2 Table. Structures and phenotypes of benzodiazepines screened against *S. mansoni.*** Data from the primary screen shown in Figure 2A. SMILES IDs are provided for all compounds screened, and phenotypes are shown for worms exposed to 30 µM test compound overnight. Blue highlighted rows indicate hits at 30 µM, red highlighted rows indicate hits at 10 µM.

