## Supplementary figures and images for "Non-sedating benzodiazepines cause contractile paralysis and tissue damage in the parasitic blood fluke *Schistosoma mansoni*"

### Supplemental Figure 1

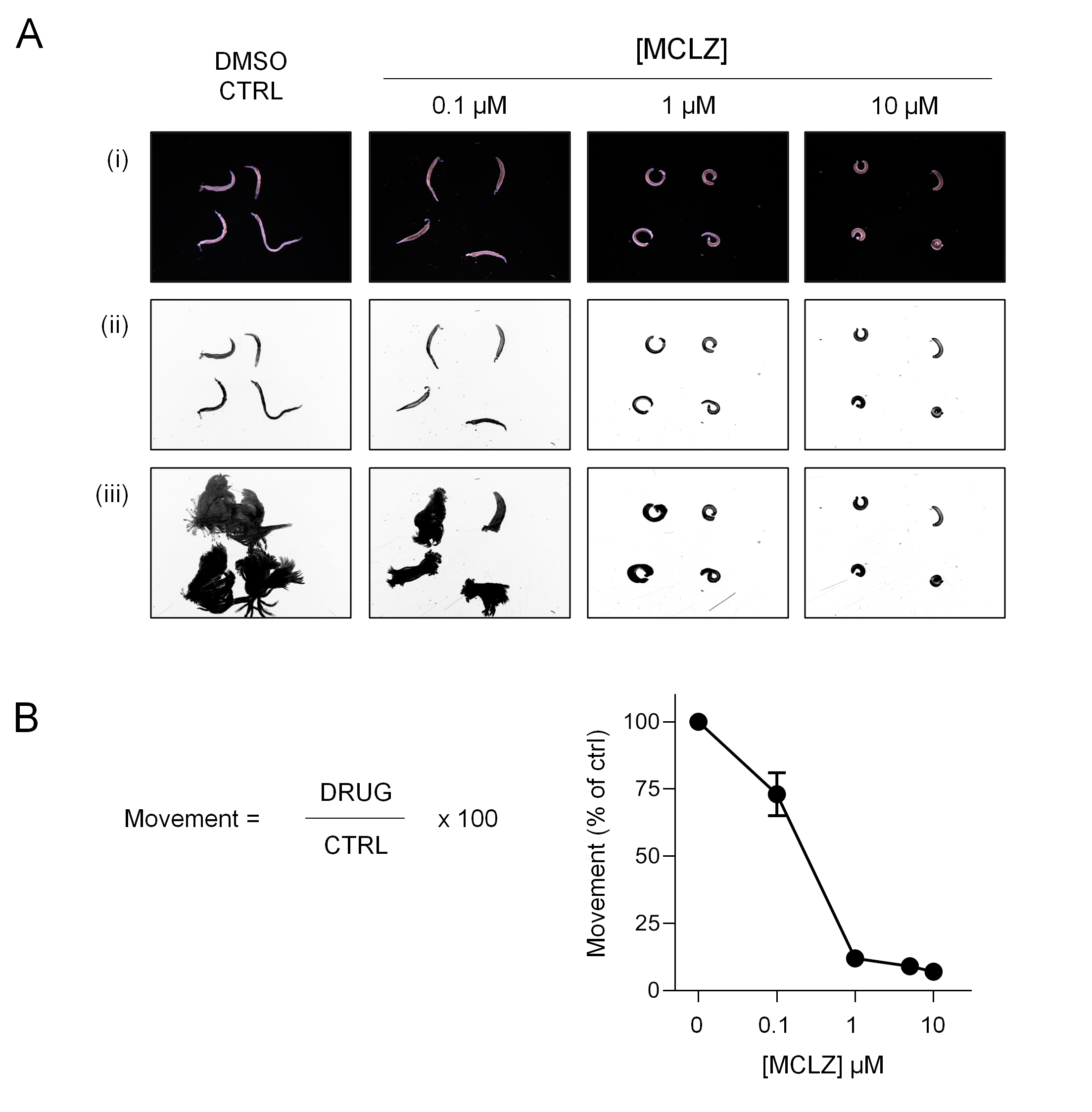
